# Evolution of insecticide resistance via parallel amino acid substitutions in insect pests and their parasitoid wasps

**DOI:** 10.1101/2022.08.18.504364

**Authors:** Lei Guo, Jia Huang

## Abstract

Insecticides have become the primary selective force in many insect species; however, whether beneficial insects developed resistance remains unknown. We analyzed the sequences of hymenopteran GABAA receptor subunit gene *Rdl* (*resistance to dieldrin*), which encodes the target of cyclodiene and phenylpyrazole insecticides. The resistance-conferring A2 S mutations were found in seven parasitoid wasps and similar amino acid replacements at homologous sites have been identified in four of their resistant hosts. Our findings indicate how parallel molecular evolution at a single amino acid site confers adaptation in both insects and their natural enemies, which may shape species interactions and community structure.

---

Pesticide resistance brings a major challenge for the sustainable control of pests, it meanwhile provides an important model to study rapid evolution under strong selective pressures, which can be used to address fundamental questions in ecology and evolutionary biology. Pesticides may also cause detrimental effects on nontarget organisms, for instance, the severe sublethal impacts of neonicotinoids on wild and managed bees have led to heavy restrictions on the use of these insecticides in Europe (Cressey 2017). Interestingly, a nontarget aquatic crustacean developed resistance to pyrethroids via mutations in the voltage-gated sodium channel, the molecular target of pyrethroids (Weston et al. 2013). However, it is largely unknown whether nontarget species, especially beneficial insects which are recurrently exposed to pesticides, evolve resistance adaptations.

The single point mutation in the GABA_A_ receptor subunit gene, *Rdl* (*resistance to dieldrin*), represents a model system for studying target site–mediated resistance to insecticides (Ffrench-Constant et al. 2000). Cyclodiene resistance is historically widespread and accounted for >60% of reported resistance cases in the 1980s, following the use of cyclodiene insecticides, including dieldrin, which was widely used during the 1950s to early 1970s (Georghiou 1986). The molecular target *Rdl* was first discovered in *Drosophila melanogaster* because a point mutation, replacing alanine with serine (A2□S, index number for M2 membrane-spanning region), of this gene confers 4,000-fold resistance to dieldrin (Ffrench-Constant et al. 1991; Ffrench-Constant et al. 1993). The mutations at position 2□ were subsequently identified from many other cyclodiene-resistant insect species (Feyereisen et al. 2015). Later, phenylpyrazole insecticides like fipronil with the same mode of action have been used to control pests starting in the 1990s. These persistent selective pressures lead to a question: have similar resistance mechanisms evolved in beneficial insects? Therefore, we analyzed the sequences of *Rdl* in Hymenoptera, because Hymenoptera, which contains pollinators, predators, and parasitoids, is not only a critical topic in ecosystem and agriculture, but also shows high sensitivity to cyclodiene and phenylpyrazole insecticides.

We focused on the M2 sequences of RDL using available genomes and transcriptomes from 59 species spanning 30 families. The A2□S mutation is found in seven parasitoids from 37 examined species (fig. 1 and supplementary fig. S1 and table S1), which is known to confer resistance to cyclodiene and phenylpyrazole insecticides via a target-site-insensitivity mechanism (Ffrench-Constant et al. 1993; Chen et al. 2006; Feyereisen et al. 2015).

**Fig. 1.**
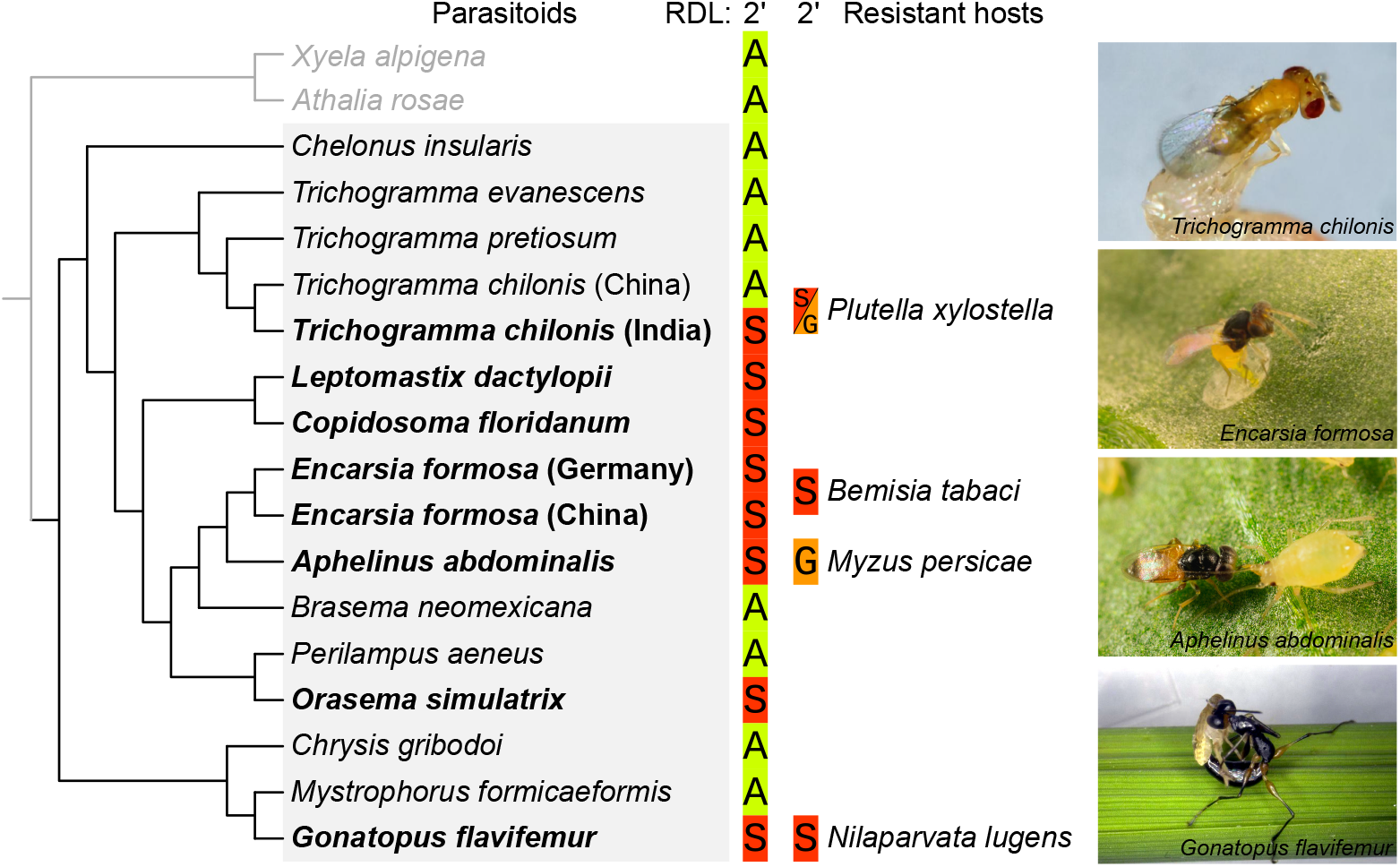
Amino acid substitutions of RDL in representative species (supplementary fig. S1 for all examined species). Gray color mark parasitoid species. The names of parasitoid species with 2□ substitutions are in bold and their respective representative host species with 2□ substitutions are shown. Only the amino acid at position 2□ is shown: green = sensitive; red and yellow = resistance. Images on the right show parasitoid wasps attacking their hosts. *Encarsia formosa* and *Aphelinus abdominalis* photos courtesy of Koppert. *Gonatopus flavifemur* image courtesy of Jiachun He (China National Rice Research Institute).

*Trichogramma* egg parasitoids are widely used for the biological control of lepidopteran pests such as the diamondback moth, *Plutella xylostella* (supplementary table S2). We observed that the *Trichogramma chilonis* strain from India has the substitution A2□S (fig. 1). The A2□S and A2□G mutations also evolved in *P. xylostella*, which showed resistance to cyclodienes and fipronil (Feyereisen et al. 2015; Wang et al. 2016). Notably, A2 S is not observed in RDL of any other *Trichogramma*, including *T. pretiosum* and *T. evanescens* (fig. 1), suggesting that the substitution evolved in recent times in *T. chilonis*. Furthermore, we collected *T. chilonis* strains from three locations in China (Beijing, Henan, and Jilin provinces) and sequenced them individually. Sequencing results showed that these strains have no amino acid substitution at position 2□. Taken together, these results suggest that point mutation in RDL of *T. chilonis* in India is an adaptation to strong insecticide selective pressure.

Dryinidae family wasps are both parasitoids and predators of Auchenorrhyncha (Hemiptera), in which the main host of *Gonatopus flavifemur* is the brown planthopper *Nilaparvata lugens*, the most serious rice pest worldwide (supplementary table S2). We observed that *Rdl* in the genome of *G. flavifemur* also encodes replacement A2□S, which was not observed in *Mystrophorus formicaeformis*, another species of the Dryinidae family (fig. 1). Then, we sequenced a field population of *G. flavifemur* collected from Zhejiang province, and confirmed the A2□S mutations. Importantly, A2 S also evolved in the brown planthopper and confers resistance to fipronil (Zhang et al. 2016). These results imply that *G. flavifemur* resists insecticides through the same point mutation found in its host.

Among the family Aphelinidae, *Encarsia formosa* is a well-known parasitoid of whiteflies and has been used as a biological-control agent since the 1920s; and *Aphelinus abdominalis* is also used for control of several aphid species (supplementary table S2). We observed that the A2□S also evolved in the RDLs of these two species (fig. 1). Since the sequenced populations of *E. Formosa* were from Germany, we collected and sequenced the strains from five locations in China (Jilin, Liaoning, Beijing, Shandong, and Zhejiang provinces), and our results confirmed that all populations have the substitution A2□S. Strikingly, the A2□S and A2□G mutations evolved in the sweet potato whitefly *Bemisia tabaci* and the green peach aphid *Myzus persicae*, respectively, which all can confer high resistance to cyclodienes (Feyereisen et al. 2015). Thus, these results suggest that the substitutions in RDL cause insensitivity to cyclodiene insecticides in *E. formosa* and *A. abdominalis* and facilitate their adaptation to insecticides.

The substitution A2□S also evolved in RDLs of *Orasema simulatrix, Leptomastix dactylopii*, and *Copidosoma floridanum* (fig. 1), which parasitize ants, citrus mealybugs, and moths in the subfamily Plusiinae, respectively (supplementary table S2). Although mutations were not reported in their host species, dieldrin and fipronil were widely used as pesticides for corn, cotton, vegetable, and citrus crops, and employed for termite and ant control, implying A2□S mutations may occur in these host species.

In summary, our findings provide the first evidence that distantly related parasitoid wasps have evolved A2□S mutations to resist cyclodiene and phenylpyrazole insecticides. Furthermore, parallel amino acid substitutions at the homologous site of RDL were found in four host-parasitoid pairs, indicating that the molecular adaptations to pesticides may reach the third trophic level. A previous field study found that non-parasitized larvae of *Manduca sexta* contain higher insecticides residues than that parasitized larvae (Dhammi 2010). Thus, the A2 S mutation-bearing pests may ingest more insecticides than that in wild-type populations and impair parasitism. As a countermeasure, their specialized parasitoids evolved the same point mutations to adapt to the otherwise lethal dose of insecticides. Our results also suggest that *Rdl* may serve as a keystone molecular marker for monitoring the effects of insecticides on beneficial insects and other nontarget animals. Finally, the widespread mutation in a commercial successful parasitoid *E. Formosa* indicates that genome engineered natural enemies with resistance-conferring mutations could be an effective method in integrated pest management.

## Materials and methods

### Identification of *Rdl* genes and phylogenetic analyses

To identify *Rdl* genes in Hymenoptera, we performed a two-step analysis: 1) we used *Drosophila melanogaster* and *Apis mellifera* genes as queries to perform BLASTp and TBLASTn search against genomes and transcriptomes, respectively; 2) we verified the candidate genes by BLASTp again without a limit of species as previously described (Guo et al. 2020; Guo et al. 2021). We took all the candidate genes that were reciprocal best hits with the *D. melanogaster Rdl* gene. Phylogenetic relationships of species were established based on previously published sources (Munro et al. 2011; Sharanowski et al. 2011; Peters et al. 2017; Bossert et al. 2019).

### Parasitoid wasps

*G. flavifemur* was provided by Dr. Qiang Fu (China National Rice Research Institute), which was collected from rice fields at Hangzhou, Zhejiang province. *T. chilonis* was provided by Dr. Liansheng Zang (Jilin Agricultural University), Kuoye Biology (http://www.kuoye.com/) (Beijing), and Henan Jiyuan Baiyun Industry Co., Ltd. (http://www.keyunnpv.cn/) (Henan province). *E. Formosa* was provided by Dr. Yinquan Liu (Zhejiang University), Dr. Liansheng Zang, Dr. Junbo Luan (Shenyang Agricultural University), Kuoye Biology, and Shandong Lubao Technology Development Co., Ltd. (http://www.saas-birc.com/) (Shandong province). *T. chilonis* and *E. Formosa* were maintained in the laboratory.

### Genotyping

Genomic DNA was extracted from a single parasitoid using the FastPure Cell/Tissue DNA Isolation Mini Kit (Vazyme Cas#DC102-01) according to the manufacturer’s protocol. Then, genomic DNA (1 μL) from the reaction was used as the PCR template for a 25 μl reaction for 35 cycles. The PCR primers spanned an approximately 200-bp region encompassing the M2 sequences (supplementary table S3). Lastly, the PCR products have checked the size using electrophoresis on a 1.2% agarose gel prior to being sequenced.

## Acknowledgments

We thank Qiang Fu (China National Rice Research Institute), Liansheng Zang (Jilin Agricultural University), Yinquan Liu (Zhejiang University), Junbo Luan (Shenyang Agricultural University) for providing the parasitoid species. This work was supported by the Zhejiang Provincial Natural Science Foundation of China (LR19C140002) and National Natural Science Foundation of China (32072496).

## Author Contributions

L.G. and J.H. designed the study, L.G. performed experiments, and L.G. and J.H. wrote the paper.

## Notes

### Competing Interest Statement

The authors have declared no competing interest.

## References

Bossert S, Murray EA, Almeida EAB, Brady SG, Blaimer BB, Danforth BN. 2019. Combining transcriptomes and ultraconserved elements to illuminate the phylogeny of Apidae. Mol. Phylogenet. Evol. 130:121–131.

Chen L, Durkin KA, Casida JE. 2006. Structural model for γ-aminobutyric acid receptor noncompetitive antagonist binding: widely diverse structure fit the same site. Proc. Natl. Acad. Sci. U. S. A. 103:5185–5190.

Cressey D. 2017. Neonics vs bees. Nature 551:156–158.

Dhammi A. 2010. Effect of Imidacloprid on Cotesia congregata, an endoparasitoid of Manduca sexta, and its translocation from host to endoparasitoid.

Feyereisen R, Dermauw W, Van Leeuwen T. 2015. Genotype to phenotype, the molecular and physiological dimensions of resistance in arthropods. Pestic. Biochem. Physiol. 121:61–77.

Ffrench-Constant RH, Anthony N, Aronstein K, Rocheleau T, Stilwell G. 2000. Cyclodiene insecticide resistance: from molecular to population genetics. Annu. Rev. Entomol. 45:449–466.

Ffrench-Constant RH, Mortlock DP, Shaffer CD, MacIntyre RJ, Roush RT. 1991. Molecular cloning and transformation of cyclodiene resistance in Drosophila: an invertebrate γ-aminobutyric acid subtype A receptor locus. Proc. Natl. Acad. Sci. U. S. A. 88:7209–7213.

Ffrench-Constant RH, Rocheleau TA, Steichen JC, Chalmers AE. 1993. A point mutation in a Drosophila GABA receptor confers insecticide resistance. Nature 363:449–451.

Georghiou GP. 1986. The magnitude of the resistance problem. In: Roush RT, Tabashnik BE, editors. Pesticide resistance: strategies and tactics for management. Washington, DC: National Academy Press. p. 14–43.

Guo L, Fan X, Qiao X, Montell C, Huang J. 2021. An octopamine receptor confers selective toxicity of amitraz on honeybees and Varroa mites. Elife 10:e68268.

Guo L, Zhou Z-D, Mao F, Fan X-Y, Liu G-Y, Huang J, Qiao X-M. 2020. Identification of potential mechanosensitive ion channels involved in texture discrimination during Drosophila suzukii egg-laying behavior. Insect Mol. Biol. 29:444–451.

Munro JB, Heraty JM, Burks RA, Hawks D, Mottern J, Cruaud A, Rasplus JY, Jansta P. 2011. A molecular phylogeny of the chalcidoidea (Hymenoptera). PLoS One 6:e27023.

Peters RS, Krogmann L, Mayer C, Donath A, Gunkel S, Meusemann K, Kozlov A, Podsiadlowski L, Petersen M, Lanfear R, et al. 2017. Evolutionary history of the Hymenoptera. Curr. Biol. 27:1013–1018.

Sharanowski BJ, Dowling APG, Sharkey MJ. 2011. Molecular phylogenetics of Braconidae (Hymenoptera: Ichneumonoidea), based on multiple nuclear genes, and implications for classification. Syst. Entomol. 36:549–572.

Wang X, Wu S, Gao W, Wu Y. 2016. Dominant inheritance of field-evolved resistance to fipronil in Plutella xylostella (Lepidoptera: Plutellidae). J. Econ. Entomol. 109:334–338.

Weston DP, Poynton HC, Wellborn GA, Lydy MJ, Blalock BJ, Sepulveda MS, Colbourne JK. 2013. Multiple origins of pyrethroid insecticide resistance across the species complex of a nontarget aquatic crustacean, Hyalella azteca. Proc. Natl. Acad. Sci. U. S. A. 110:16532–16537.

Zhang Y, Meng X, Yang Y, Li H, Wang X, Yang B, Zhang J, Li C, Millar NS, Liu Z. 2016. Synergistic and compensatory effects of two point mutations conferring target-site resistance to fipronil in the insect GABA receptor RDL. Sci. Rep. 6:32335.

