## Supplemental tables and figures for "Evolution of insecticide resistance via parallel amino acid substitutions in insect pests and their parasitoid wasps"

Supplementary tables

### Table S1 List of *Rdl* sequences and accession numbers used in this study (when species have evolved mutation in position 2ʹ are highlighted in yellow)

| **Order** | **Family** | **Species** | **M2 2ʹ position** | ***Rdl* Source** | **Accession Number** | **Reference** | **Data Source** |
| --- | --- | --- | --- | --- | --- | --- | --- |
| Hymenoptera | Xyelidae | *Xyela alpigena* | A | GenBank | GBVH01017944.1 | (Peters et al. 2017) | Transcriptome |
| Hymenoptera | Diprionidae | *Neodiprion lecontei* | A | GenBank | XM_015653665.1 | (Vertacnik et al. 2021) | Genome |
| Hymenoptera | Tenthredinidae | *Nematus ribesii* | A | GenBank | GBQB01018038.1 | (Peters et al. 2017) | Transcriptome |
| Hymenoptera | Tenthredinidae | *Athalia rosae* | A | GenBank | XM_012409466.3 | (Oeyen et al. 2020) | Genome |
| Hymenoptera | Cephidae | *Cephus cinctus* | A | GenBank | XM_015733587.2 | (Robertson et al. 2018) | Genome |
| Hymenoptera | Orussidae | *Orussus abietinus* | A | GenBank | XM_023433906.1 | (Oeyen et al. 2020) | Genome |
| Hymenoptera | Braconidae | *Fopius arisanus* | A | GenBank | XM_011301113.1 | (Geib et al. 2017) | Genome |
| Hymenoptera | Braconidae | *Macrocentrus cingulum* | A | InsectBase 2.0 | MCIN04062-TA | (Yin et al. 2018) | Genome |
| Hymenoptera | Braconidae | *Chelonus insularis* | A | GenBank | XM_035082937.1 | (Poelchau et al. 2015) | Genome |
| Hymenoptera | Braconidae | *Microplitis demolitor* | A | GenBank | XM_008551582.1 | (Burke et al. 2018) | Genome |
| Hymenoptera | Braconidae | *Cotesia congregata* | A | GenBank | CAD6240252.1 | (Gauthier et al. 2021) | Transcriptome |
| Hymenoptera | Braconidae | *Cotesia vestalis* | A | GenBank | GAUP02020021.1 | (Misof et al. 2014) | Transcriptome |
| Hymenoptera | Megaspilidae | *Dendrocerus carpenteri* | A | GenBank | GBOS01019186.1 | (Misof et al. 2014) | Transcriptome |
| Hymenoptera | Figitidae | *Leptopilina boulardi* | A | GenBank | GITC01127536.1 | (Peters et al. 2017) | Transcriptome |
| Hymenoptera | Trichogrammatidae | *Trichogramma evanescens* | A | GenBank | GBMC01010992.1 | (Peters et al. 2018) | Transcriptome |
| Hymenoptera | Trichogrammatidae | *Trichogramma pretiosum* | A | GenBank | XM_014371636.2 | (Lindsey et al. 2018) | Genome |
| Hymenoptera | Trichogrammatidae | *Trichogramma chilonis* | S | GenBank | GGKM01012770.1 | (Kerima et al. 2018) | Transcriptome |
| Hymenoptera | Encyrtidae | *Leptomastix dactylopii* | S | GenBank | GBNE01033321.1 | (Peters et al. 2018) | Transcriptome |
| Hymenoptera | Encyrtidae | *Copidosoma floridanum* | S | GenBank | XM_023391076.1 | (Thomas et al. 2020) | Genome |
| Hymenoptera | Aphelinidae | *Encarsia formosa* | S | GenBank | GBVN01041245.1 | (Misof et al. 2014) | Transcriptome |
| Hymenoptera | Aphelinidae | *Aphelinus abdominalis* | S | GenBank | GBTK01038292.1 | (Misof et al. 2014) | Transcriptome |
| Hymenoptera | Eupelmidae | *Brasema neomexicana* | A | GenBank | GBUE01028569.1 | (Peters et al. 2018) | Transcriptome |
| Hymenoptera | Eupelmidae | *Eupelmus urozonus* | A | GenBank | GBMN01021006.1 | (Peters et al. 2018) | Transcriptome |
| Hymenoptera | Eulophidae | *Diglyphus isaea* | A | GenBank | GBLP01023234.1 | (Peters et al. 2017) | Transcriptome |
| Hymenoptera | Eulophidae | *Tamarixia radiata* | A | GenBank | GBMM01031177.1 | (Peters et al. 2018) | Transcriptome |
| Hymenoptera | Eulophidae | *Elasmus sp.* | A | GenBank | GCUU01024471.1 | (Peters et al. 2018) | Transcriptome |
| Hymenoptera | Pteromalidae | *Pachycrepoideus vindemmiae* | A | GenBank | GBWL01025773.1 | (Peters et al. 2018) | Transcriptome |
| Hymenoptera | Pteromalidae | *Philotrypesis parca* | A | GenBank | GBOV01019554.1 | (Peters et al. 2018) | Transcriptome |
| Hymenoptera | Pteromalidae | *Nasonia giraulti* | A | GenBank | GBEC01014311.1 | (Hoedjes et al. 2015) | Transcriptome |
| Hymenoptera | Pteromalidae | *Nasonia vitripennis* | A | GenBank | XM_016990297.3 | (Benetta et al. 2020) | Genome |
| Hymenoptera | Pteromalidae | *Pteromalus puparum* | A | GenBank | GECT01028516.1 | (Yan et al. 2016) | Transcriptome |
| Hymenoptera | Chalcididae | *Brachymeria minuta* | A | GenBank | GBLE01013399.1 | (Peters et al. 2017) | Transcriptome |
| Hymenoptera | Perilampidae | *Perilampus aeneus* | A | GenBank | GBLT01022262.1 | (Peters et al. 2018) | Transcriptome |
| Hymenoptera | Eucharitidae | *Orasema simulatrix* | S | GenBank | GBTU01000582.1 | (Peters et al. 2017) | Transcriptome |
| Hymenoptera | Torymidae | *Torymus bedeguaris* | A | GenBank | GBQA01037379.1 | (Peters et al. 2017) | Transcriptome |
| Hymenoptera | Torymidae | *Bootanomyia dorsalis* | A | GenBank | GBPA01024904.1 | (Peters et al. 2018) | Transcriptome |
| Hymenoptera | Agaonidae | *Ceratosolen solmsi marchali* | A | GenBank | XM_011497468.1 | (Xiao et al. 2013) | Genome |
| Hymenoptera | Gasteruptiidae | *Gasteruption tournieri* | A | GenBank | GBVG01020305.1 | (Peters et al. 2017) | Transcriptome |
| Hymenoptera | Stephanidae | *Stephanus serrator* | A | GenBank | GBMU01009039.1 | (Peters et al. 2017) | Transcriptome |
| Hymenoptera | Chrysididae | *Chrysis gribodoi* | A | GenBank | GCUE01045579.1 | (Pauli et al. 2021) | Transcriptome |
| Hymenoptera | Dryinidae | *Mystrophorus formicaeformis* | A | GenBank | GDLQ01003233.1 | (Pauli et al. 2021) | Transcriptome |
| Hymenoptera | Dryinidae | *Gonatopus flavifemur* | S | InsectBase 2 | Gfla020038.1 | (Yang et al. 2021) | Genome |
| Hymenoptera | Vespidae | *Polistes metricus* | A | GenBank | GDHQ01058842.1 | (Berens et al. 2017) | Transcriptome |
| Hymenoptera | Vespidae | *Polistes dominula* | A | GenBank | XM_015328436.1 | (Standage et al. 2016) | Genome |
| Hymenoptera | Chyphotidae | *Chyphotes mellipes* | A | GenBank | GAXL01021678.1 | (Johnson et al. 2013) | Transcriptome |
| Hymenoptera | Tiphiidae | *Methocha articulata* | A | GenBank | GBTZ01033330.1 | (Peters et al. 2017) | Transcriptome |
| Hymenoptera | Tiphiidae | *Brachycistis timberlakei* | A | GenBank | GAZU01005585.1 | (Johnson et al. 2013) | Transcriptome |
| Hymenoptera | Formicidae | *Camponotus floridanus* | A | GenBank | XM_011253963.3 | (Shields et al. 2018) | Genome |
| Hymenoptera | Formicidae | *Harpegnathos saltator* | A | GenBank | XM_011139263.3 | (Shields et al. 2018) | Genome |
| Hymenoptera | Crabronidae | *Astata minor* | A | GenBank | GBLB01005644.1 | (Peters et al. 2017) | Transcriptome |
| Hymenoptera | Crabronidae | *Psenulus fuscipennis* | A | GenBank | GBNH01019008.1 | (Petersen et al. 2017) | Transcriptome |
| Hymenoptera | Megachilidae | *Megachile rotundata* | A | GenBank | XM_012288275. | (Kapheim et al. 2015) | Genome |
| Hymenoptera | Megachilidae | *Osmia lignaria* | A | GenBank | XM_034325711.1 | (Poelchau et al. 2015) | Genome |
| Hymenoptera | Apidae | *Eufriesea mexicana* | A | GenBank | XM_017899520.1 | (Kapheim et al. 2015) | Genome |
| Hymenoptera | Apidae | *Apis florea* | A | GenBank | XM_031915912.1 | (Fouks et al. 2021) | Genome |
| Hymenoptera | Apidae | *Apis dorsata* | A | GenBank | XM_006618939.2 | (Oppenheim et al. 2020) | Genome |
| Hymenoptera | Apidae | *Apis mellifera* | A | GenBank | XM_006565106.3 | ( The Honeybee Genome Sequencing Consortium 2006) | Genome |
| Hymenoptera | Apidae | *Apis cerana* | A | GenBank | XM_017059006.2 | (Park et al. 2015) | Genome |
| Hymenoptera | Apidae | *Bombus impatiens* | A | GenBank | XM_024365028.2 | (Sadd et al. 2015) | Genome |

### Table S2 Host preference of the parasitoid species used this study.

| **Family** | **Species** | **Host preference** | **Reference** |
| --- | --- | --- | --- |
| Braconidae | *Chelonus insularis* | *Spodoptera frugiperda* | (Roque-Romero et al. 2020) |
| Trichogrammatidae | *Trichogramma pretiosum* | *Anagasta kuehniella >* *Spodoptera frugiperda* | (Siqueira et al. 2012) |
| Trichogrammatidae | *Trichogramma evanescens* | *Mamestra brassicae* > *Pieris brassicae* ≈ *Pieris rapae* | (Marianne J.van Dijken, Marian Kole 1986) |
| Trichogrammatidae | *Trichogramma chilonis* | *Plutella xylostella* | (Wührer and Hassan 1993) |
| Encyrtidae | *Leptomastix dactylopii* | *Planococcus citri* | (Cloyd and Sadof 2000) |
| Encyrtidae | *Copidosoma floridanum* | *Trichoplusia ni* | (Lampert and Bowers 2010) |
| Aphelinidae | *Encarsia formosa* | *Bemisia tabaci* | (Hoddle et al. 1998) |
| Aphelinidae | *Aphelinus abdominalis* | *Rhopalosiphum padi* > *Myzus persicae* > *Macrosiphum euphorbiae* | (Velasco-Hernández et al. 2017) |
| Eulophidae | *Tamarixia radiata* | *Diaphorina citri* | (Hoddle and Pandey 2014) |
| Perilampidae | *Perilampus aeneus* | *Athalia rosae* | {Formatting Citation} |
| Eucharitidae | *Orasema simulatrix* | *Pheidole desertorum* | (Carey et al. 2012) |
| Chrysididae | *Chrysis gribodoi* | / | / |
| Dryinidae | *Mystrophorus formicaeformis* | / | / |
| Dryinidae | *Gonatopus flavifemur* | *Nilaparvata lugens* | (He et al. 2020) |

### Table S3 Primers used in this study

| **Target** | **Forward primer (5’-3’)** | **Reverse primer (5’-3’)** |
| --- | --- | --- |
| *Gonatopus flavifemur Rdl* | CCATCGGGACTGATCGTCAT | AACAGCGAGGCGAAAACCAT |
| *Trichogramma chilonis Rdl* | CGTCAGGACTGATCGTCATCATA | AGCAACGATGCGAAGACCA |
| *Encarsia formosa Rdl* | TGGGCTACTACCTGATCCAGA | AAACAGGTGCCCAGGTAGAC |

Supplementary figure

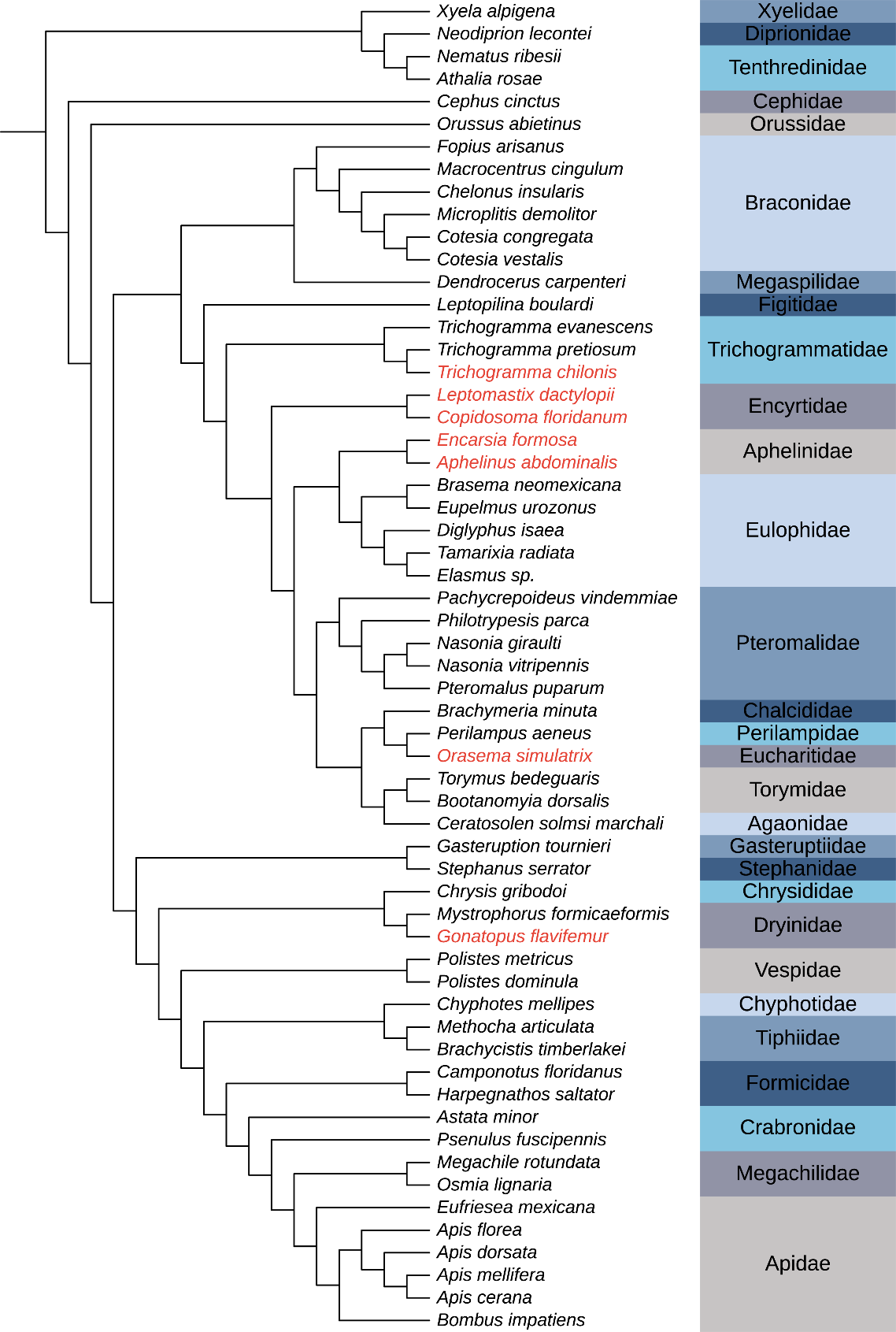

### Fig. S1 Phylogeny showing relationships of the sampled species.

The names of parasitoid wasp species with amino acid substitutions are in red. Phylogenetic relationships of species were established based on previously published sources (see methods).

Supplementary references

Benetta ED, Antoshechkin I, Yang T, My Nguyen HQ, Ferree PM, Akbari OS. 2020. Genome elimination mediated by gene expression from a selfish chromosome. *Sci. Adv.* 6:eaaz9808.

Berens AJ, Tibbetts EA, Toth AL. 2017. Cognitive specialization for learning faces is associated with shifts in the brain transcriptome of a social wasp. *J. Exp. Biol.* 220:2149–2153.

Burke GR, Walden KKO, Whitfield JB, Robertson HM, Strand MR. 2018. Whole genome sequence of the parasitoid wasp microplitis demolitor that harbors an endogenous virus mutualist. *G3 Genes, Genomes, Genet.* 8:2875–2880.

CABI. 2022. *Perilampus aeneus*. In: Invasive species compendium. Wallingford, UK: CAB International. www.cabi.org/isc.

Carey B, Visscher K, Heraty J. 2012. Nectary use for gaining access to an ant host by the parasitoid *Orasema simulatrix* (Hymenoptera, Eucharitidae). *J. Hymenopt. Res.* 27:47–65.

Cloyd RA, Sadof CS. 2000. Effects of plant architecture on the attack rate of *Leptomastix dactylopii* (Hymenoptera: Encyrtidae), a parasitoid of the citrus mealybug (Homoptera: Pseudococcidae). *Environ. Entomol.* 29:535–541.

Fouks B, Brand P, Nguyen HN, Herman J, Camara F, Ence D, Hagen DE, Hoff KJ, Nachweide S, Romoth L, et al. 2021. The genomic basis of evolutionary differentiation among honey bees. *Genome Res.* 31:1203–1215.

Gauthier J, Boulain H, van Vugt JJFA, Baudry L, Persyn E, Aury JM, Noel B, Bretaudeau A, Legeai F, Warris S, et al. 2021. Chromosomal scale assembly of parasitic wasp genome reveals symbiotic virus colonization. *Commun. Biol.* 4:104.

Geib SM, Liang GH, Murphy TD, Sim SB. 2017. Whole genome sequencing of the braconid parasitoid wasp *Fopius arisanus*, an important biocontrol agent of pest tepritid fruit flies. *G3 Genes, Genomes, Genet.* 7:2407–2411.

He J, He Y, Lai F, Chen X, Fu Q. 2020. Biological traits of the pincer wasp *Gonatopus flavifemur* (Esaki & Hashimoto) associated with different stages of its host, the brown planthopper, *Nilaparvata lugens* (Stål). *Insects* 11:279.

Hoddle MS, Van Driesche RG, Sanderson JP. 1998. Biology and use of the whitefly parasitoid *Encarsia formosa*. *Annu. Rev. Entomol.* 43:645–669.

Hoddle MS, Pandey R. 2014. Host range testing of *Tamarixia radiata* (Hymenoptera: Eulophidae) sourced from the Punjab of Pakistan for classical biological control of *Diaphorina citri* (Hemiptera: Liviidae: Euphyllurinae: Diaphorinini) in California. *J. Econ. Entomol.* 107:125–136.

Hoedjes KM, Smid HM, Schijlen EGWM, Vet LEM, van Vugt JJFA. 2015. Learning-induced gene expression in the heads of two Nasonia species that differ in long-term memory formation. *BMC Genomics* [Internet] 16:162.

Johnson BR, Borowiec ML, Chiu JC, Lee EK, Atallah J, Ward PS. 2013. Phylogenomics resolves evolutionary relationships among ants, bees, and wasps. *Curr. Biol.* 23:2058–2062.

Kapheim KM, Pan H, Li C, Salzberg SL, Puiu D, Magoc T, Robertson HM, Hudson ME, Venkat A. 2015. Genomic signatures of evolutionary transitions from solitary to group living. *Science* 348:1139–1144.

Kerima OZ, Niranjana P, Vinay Kumar BS, Ramachandrappa R, Puttappa S, Lalitha Y, Jalali SK, Ballal CR, Thulasiram H V. 2018. De novo transcriptome analysis of the egg parasitoid *Trichogramma chilonis* Ishii (Hymenoptera: Trichogrammatidae): a biological control agent. *Gene Reports* 13:115–129.

Lampert EC, Bowers MD. 2010. Host plant species affects the quality of the generalist *Trichoplusia ni* as a host for the polyembryonic parasitoid Copidosoma floridanum. *Entomol. Exp. Appl.* 134:287–295.

Lindsey ARI, Kelkar YD, Wu X, Sun D, Martinson EO, Yan Z, Rugman-Jones PF, Hughes DST, Murali SC, Qu J, et al. 2018. Comparative genomics of the miniature wasp and pest control agent *Trichogramma pretiosum*. *BMC Biol.* 16:54.

Marianne J.van Dijken, Marian Kole JC va. L and AMB. 1986. Host-preference studies with *Trichogramma evanescens* Westwood (Hym., Trichogrammatidae) for *Mamestra brassicae*, *Pieris brassicae* and *Pieris rapae*. *J. Appl. Entomol.* 101:64–85.

Misof B, Liu S, Meusemann K, Peters RS, Donath A, Mayer C, Frandsen PB, Ware J, Flouri T, Beutel RG, et al. 2014. Phylogenomics resolves the timing and pattern of insect evolution. *Science* 346:763–767.

Oeyen JP, Baa-Puyoulet P, Benoit JB, Beukeboom LW, Bornberg-Bauer E, Buttstedt A, Calevro F, Cash EI, Chao H, Charles H, et al. 2020. Sawfly genomes reveal evolutionary acquisitions that fostered the mega-radiation of parasitoid and eusocial hymenoptera. *Genome Biol. Evol.* 12:1099–1188.

Oppenheim S, Cao X, Rueppel O, Krongdang S, Phokasem P, Desalle R, Goodwin S, Xing J, Chantawannakul P, Rosenfeld JA. 2020. Whole genome sequencing and assembly of the Asian honey bee *Apis dorsata*. *Genome Biol. Evol.* 12:3677–3683.

Park D, Jung JW, Choi B-S, Jayakodi M, Lee J, Lim J, Yu Y, Choi Y-S, Lee M-L, Park Y, et al. 2015. Uncovering the novel characteristics of Asian honey bee, *Apis cerana*, by whole genome sequencing. *BMC Genomics* 16:1.

Pauli T, Meusemann K, Kukowka S, Sann M, Donath A, Mayer C, Oeyen JP, Ballesteros Y, Berg A, Van Den Berghe E, et al. 2021. Analysis of RNA-Seq, DNA target enrichment, and sanger nucleotide sequence data resolves deep splits in the phylogeny of cuckoo wasps (Hymenoptera: Chrysididae). *Insect Syst. Divers.* 5:1.

Peters RS, Krogmann L, Mayer C, Donath A, Gunkel S, Meusemann K, Kozlov A, Podsiadlowski L, Petersen M, Lanfear R, et al. 2017. Evolutionary history of the Hymenoptera. *Curr. Biol.* 27:1013–1018.

Peters RS, Niehuis O, Gunkel S, Bläser M, Mayer C, Podsiadlowski L, Kozlov A, Donath A, van Noort S, Liu S, et al. 2018. Transcriptome sequence-based phylogeny of chalcidoid wasps (Hymenoptera: Chalcidoidea) reveals a history of rapid radiations, convergence, and evolutionary success. *Mol. Phylogenet. Evol.* 120:286–296.

Petersen M, Meusemann K, Donath A, Dowling D, Liu S, Peters RS, Podsiadlowski L, Vasilikopoulos A, Zhou X, Misof B, et al. 2017. Orthograph: a versatile tool for mapping coding nucleotide sequences to clusters of orthologous genes. *BMC Bioinformatics* 18:111.

Poelchau M, Childers C, Moore G, Tsavatapalli V, Evans J, Lee CY, Lin H, Lin JW, Hackett K. 2015. The i5k Workspace@NAL-enabling genomic data access, visualization and curation of arthropod genomes. *Nucleic Acids Res.* 43:D714–D719.

Robertson HM, Waterhouse RM, Walden KKO, Ruzzante L, Reijnders MJMF, Coates BS, Legeai F, Gress JC, Biyiklioglu S, Weaver DK, et al. 2018. Genome sequence of the wheat stem sawfly, Cephus cinctus, representing an early-branching lineage of the hymenoptera, illuminates evolution of hymenopteran chemoreceptors. *Genome Biol. Evol.* 10:2997–3011.

Roque-Romero L, Cisneros J, Rojas JC, Ortiz-Carreon FR, Malo EA. 2020. Attraction of *Chelonus insularis* to host and host habitat volatiles during the search of *Spodoptera frugiperda* eggs. *Biol. Control.* 140:104100.

Sadd BM, Barribeau SM, Bloch G, de Graaf DC, Dearden P, Elsik CG, Gadau J, Grimmelikhuijzen CJP, Hasselmann M, Lozier JD, et al. 2015. The genomes of two key bumblebee species with primitive eusocial organization. *Genome Biol.* 16:76.

Shields EJ, Sheng L, Weiner AK, Garcia BA, Bonasio R. 2018. High-quality genome assemblies reveal long non-coding RNAs expressed in ant brains. *Cell Rep.* 23:3078–3090.

Siqueira JR, de Freitas Bueno RCO, De Freitas Bueno A, Vieira SS. 2012. Host preference of the egg parasitoid *Trichogramma pretiosum*. *Cienc. Rural* 42:1–5.

Standage DS, Berens AJ, Glastad KM, Severin AJ, Brendel VP, Toth AL. 2016. Genome, transcriptome and methylome sequencing of a primitively eusocial wasp reveal a greatly reduced DNA methylation system in a social insect. *Mol. Ecol.* 25:1769–1784.

The Honeybee Genome Sequencing Consortium. 2006. Insights into social insects from the genome of the honeybee *Apis mellifera*. *Nature* 443:931–949.

Thomas GWC, Dohmen E, Hughes DST, Murali SC, Poelchau M, Glastad K, Anstead CA, Ayoub NA, Batterham P, Bellair M, et al. 2020. Gene content evolution in the arthropods. *Genome Biol.* 21:15.

Velasco-Hernández MC, Desneux N, Ramírez-Martínez MM, Cicero L, Ramirez-Romero R. 2017. Host species suitability and instar preference of *Aphidius ervi* and *Aphelinus abdominalis.* *Entomol. Gen.* 36:347–367.

Vertacnik KL, Herrig DK, Godfrey RK, Hill T, Geib SM, Unckless RL, Nelson DR, Linnen CR. 2021. Ecological correlates of gene family size in a pine-feeding sawfly genome and across Hymenoptera. *bioRxiv*.

Wührer BG, Hassan SA. 1993. Selection of effective species/strains of Trichogramma (Hym., Trichogrammatidae) to control the diamondback moth *Plutella xylostella* L. (Lep., Plutellidae). *J. Appl. Entomol.* 116:80–89.

Xiao JH, Yue Z, Jia LY, Yang XH, Niu LH, Wang Z, Zhang P, Sun BF, He SM, Li Z, et al. 2013. Obligate mutualism within a host drives the extreme specialization of a fig wasp genome. *Genome Biol.* 14:R141.

Yan Z, Fang Q, Wang L, Liu J, Zhu Y, Wang F, Li F, Werren JH, Ye G. 2016. Insights into the venom composition and evolution of an endoparasitoid wasp by combining proteomic and transcriptomic analyses. *Sci. Rep.* 6:19604.

Yang Y, Ye X, Dang C, Cao Y, Hong R, Sun YH, Xiao S, Mei Y, Xu L, Fang Q, et al. 2021. Genome of the pincer wasp *Gonatopus flavifemur* reveals unique venom evolution and a dual adaptation to parasitism and predation. *BMC Biol.* 19:1–24.

Yin C, Li M, Hu J, Lang K, Chen Q, Liu J, Guo D, He K, Dong Y, Luo J, et al. 2018. The genomic features of parasitism, polyembryony and immune evasion in the endoparasitic wasp *Macrocentrus cingulum*. *BMC Genomics* 19:420.
